# Self-Interaction Nanoparticle Spectroscopy Predicts High-Concentration Viscosity of Therapeutic IgG1 Antibodies

**DOI:** 10.64898/2026.04.16.719068

**Authors:** Santosh Kumar Paidi, Julissa Ibrahim, Kateryna Stepurska, Jonathan Zarzar, Saeed Izadi, Erina Rude, Steven Luu, Daniel Kovner, Kelly O’Connor, Karenna Bol, Shrenik Mehta, Nisana Andersen, Nicole Stephens, Emily Makowski, Joel Heisler, Trevor Swartz, Paul J. Carter, Tomasz Baginski

## Abstract

Predicting high-concentration viscosity of monoclonal antibodies such as IgG1 is crucial for their development as therapeutics for subcutaneous delivery. Unfortunately, traditional experimental rheometry methods for assessing viscosity are low-throughput. This study evaluates Self-Interaction Nanoparticle Spectroscopy (SINS) assays—specifically charge-stabilized SINS (CS-SINS) and PEG-stabilized SINS (PS-SINS)—for high-throughput viscosity prediction. We characterized 96 IgG_1_ antibodies, assessing SINS against *in silico* descriptors and dynamic light scattering (DLS) data. CS-SINS showed strong correlation with charge, offering limited additional utility. In contrast, PS-SINS provided orthogonal information; integrating it with *in silico* data and DLS significantly improved random forest model accuracy for binary viscosity classification. PS-SINS measurements in multiple buffers captured complementary information, achieving comparable accuracy without DLS. Importantly, PS-SINS scores exhibited a strong logarithmic relationship (r=0.98) with high-concentration viscosity in Fc variants of clinical antibodies, suggesting a direct mechanistic link. Furthermore, PS-SINS performed reliably with one column purified (protein A) samples, supporting its early-stage application. These findings establish PS-SINS as a high-throughput tool to accelerate the developability assessment of antibody candidates.

## 1. Introduction

The current era of biopharmaceutical development is characterized by a significant emphasis on monoclonal antibody (mAb) therapeutics and their subcutaneous delivery.^1,2^ These molecules have transformed therapeutic strategies for a wide spectrum of diseases, including multiple cancers and autoimmune disorders.^3^ Predicting the high-concentration viscosity and developability of monoclonal antibody (mAb) therapeutics is critical for successful pharmaceutical development.^4,5^ High viscosity poses significant challenges in manufacturing, formulation, and administration, potentially impacting drug efficacy and patient experience.^6,7^ Traditional cone-and-plate rheometry for assessing high-concentration viscosity often require substantial quantities of protein (≥30 mg) and time, hindering application in early-stage screening.^8,9^ While rheological measurements provide direct viscosity values, and DLS measures diffusion interaction, both are low-throughput and unsuitable for early screening of large number of molecules.^10^

Emerging Self-Interaction Nanoparticle Spectroscopy (SINS) techniques present a promising alternative for rapid evaluation of developability liabilities of antibodies by capturing them on gold nanoparticles and facilitating the interactions that dictate their propensity for self-association, which is an important factor influencing high-concentration viscosity.^11–14^ The optical readout of the surface plasmon shift of the gold nanoparticles caused by test antibody-mediated changes in their interparticle distance enables high-throughput detection during early development with limited antibody samples. Originally developed affinity-capture self-interaction nanoparticle spectroscopy (AC-SINS) is widely used for early antibody drug candidate screening to identify high-concentration liabilities, but is unsuitable for all formulation conditions, particularly histidine-based buffers.^11,12,14^ The next generation assay, charge-stabilized SINS (CS-SINS) leveraged poly-lysine, a positively charged polymer, to stabilize the antibody–gold conjugates.^13^ Around the same time, PEG-stabilized SINS (PS-SINS) was reported which used inert polyethylene glycol (PEG) to stabilize the antibody-gold conjugates and demonstrated the suitability of the method to evaluate the developability liability of antibody molecules in multiple buffers.^14^ However, these groups neither quantitatively characterize their ability to predict viscosity as a binary classification task nor compared the readouts of these assays with *in silico* predictors of charge and hydrophobicity attributes of molecules. Furthermore, there is also a need to assess the direct relationship between viscosity and PS-SINS behavior in a controlled set of antibody mutants with varying viscosity levels.

In this manuscript, through *in silico* and experimental characterization of a diverse set of 96 IgG_1_ molecules that span a wide range of high-concentration viscosity, charge and hydrophobicity, we present a comprehensive assessment of the relevance of CS-SINS and PS-SINS assays for predicting high-concentration viscosity (∼180 mg/mL, in 20 mM histidine acetate pH 5.8 buffer) of antibody therapeutics (**Figure 1**). We automated the plate setup part of sample preparation of both the assays using a liquid handler for high-throughput experimentation. We compared the performance of CS-SINS and PS-SINS assays for predicting high-concentration viscosity. By combining *in silico* molecular descriptors with experimental data from SINS assays and dynamic light scattering (DLS), we trained supervised random forest-based binary classifiers for prediction of high-concentration viscosity. We evaluated the predictive power of PS-SINS measurements obtained in multiple buffers (phosphate-buffered saline pH 7.4 (PBS), 10 mM histidine hydrochloride pH 6.0, 20 mM histidine acetate pH 5.8, 200 mM arginine succinate pH 5.8, and 200 mM arginine chloride pH 5.8) for viscosity prediction. We also leveraged Fc mutation variants of two clinical antibodies that respectively exhibit low and high viscosities at high-concentration (180 mg/mL, in 20 mM histidine acetate pH 5.8 buffer) to examine the direct relationship between PS-SINS score and high-concentration viscosity. Finally, we performed PS-SINS on the one-step purified (protein A chromatography) samples of the same set of 96 antibodies to assess the impact of antibody purity on the PS-SINS assay performance. Taken together, the results of this study are expected to help early molecule development teams select therapeutic antibody candidates with favorable high-concentration properties.

**Figure 1.**
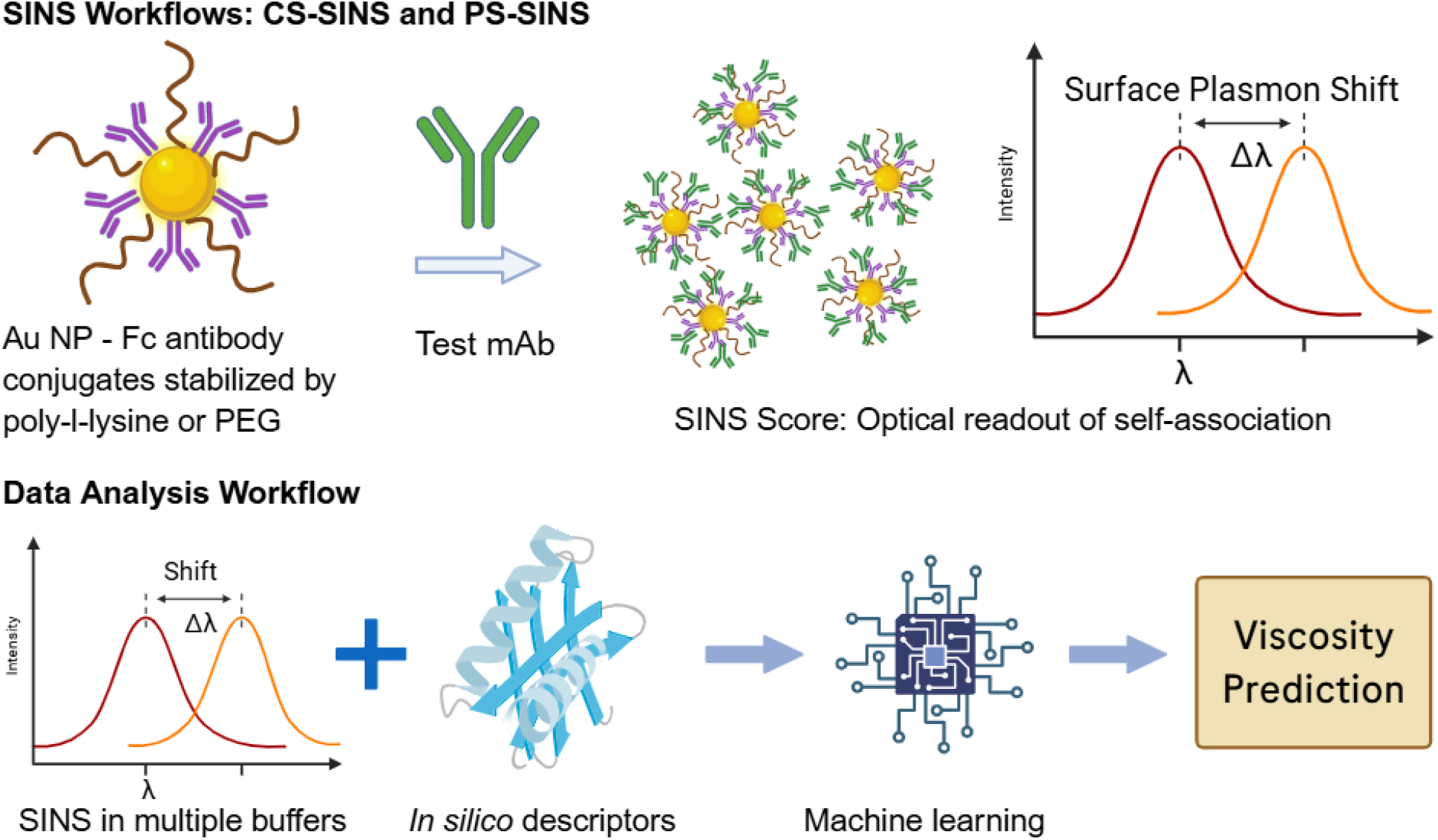
SINS for prediction of high concentration viscosity. The simplified schematic of CS-SINS, PS-SINS, and data analysis workflows is shown. The test antibodies are incubated with capture antibody conjugated nanoparticles and shift in the surface plasmon wavelength of the gold nanoparticles due to self-association is measured. The SINS results and *in silico* molecular descriptors are leveraged for building classification models for high-concentration viscosity prediction.

## 2. Results

### 2.1. CS-SINS strongly correlates with charge and provides limited additional utility in comparison to *in silico* charge descriptors

A comprehensive suite of Fab-centric *in silico* molecular descriptors, encompassing both charge and hydrophobicity properties, were computed for each antibody to characterize their behavior. These descriptors were largely derived using the Adaptive Poisson-Boltzmann Solver (APBS) for electrostatic potential^15^ and the Spatial Aggregation Propensity (SAP) method for hydrophobicity^16^, with many also incorporating insights from Gaussian accelerated molecular dynamics (GaMD)^17^ and Molecular Operating Environment (MOE) simulations.^18^

We examined the correlation between *in silico* molecular descriptors of the molecules and experimentally derived measures of viscosity, DLS-derived diffusion interaction parameter (*k*_D_), and SINS scores (PS-SINS and CS-SINS scores in 10 mM histidine hydrochloride pH 6.0 buffer unless noted otherwise) as a heatmap (**Figure 2A**). The CS-SINS scores showed strong negative correlation (Pearson correlation coefficient, r = -0.70) with charge-related molecular descriptors including isoelectric point (pI) (**Figure 2A-B**). PS-SINS scores, on the other hand, did not show strong correlation with either charge or hydrophobicity descriptors used in this study. A scatterplot between CS-SINS and PS-SINS scores showed that while both the scores correlated with each other for the common calibration standard molecules used in both the assays for score normalization, they did not show significant correlation across the entire dataset (**Figure 2C**). Both CS-SINS and PS-SINS scores show weak univariate correlations with viscosity with PS-SINS (r = 0.29) slightly outperforming CS-SINS (r = 0.17) (**Figure 2D-E**).

**Figure 2.**
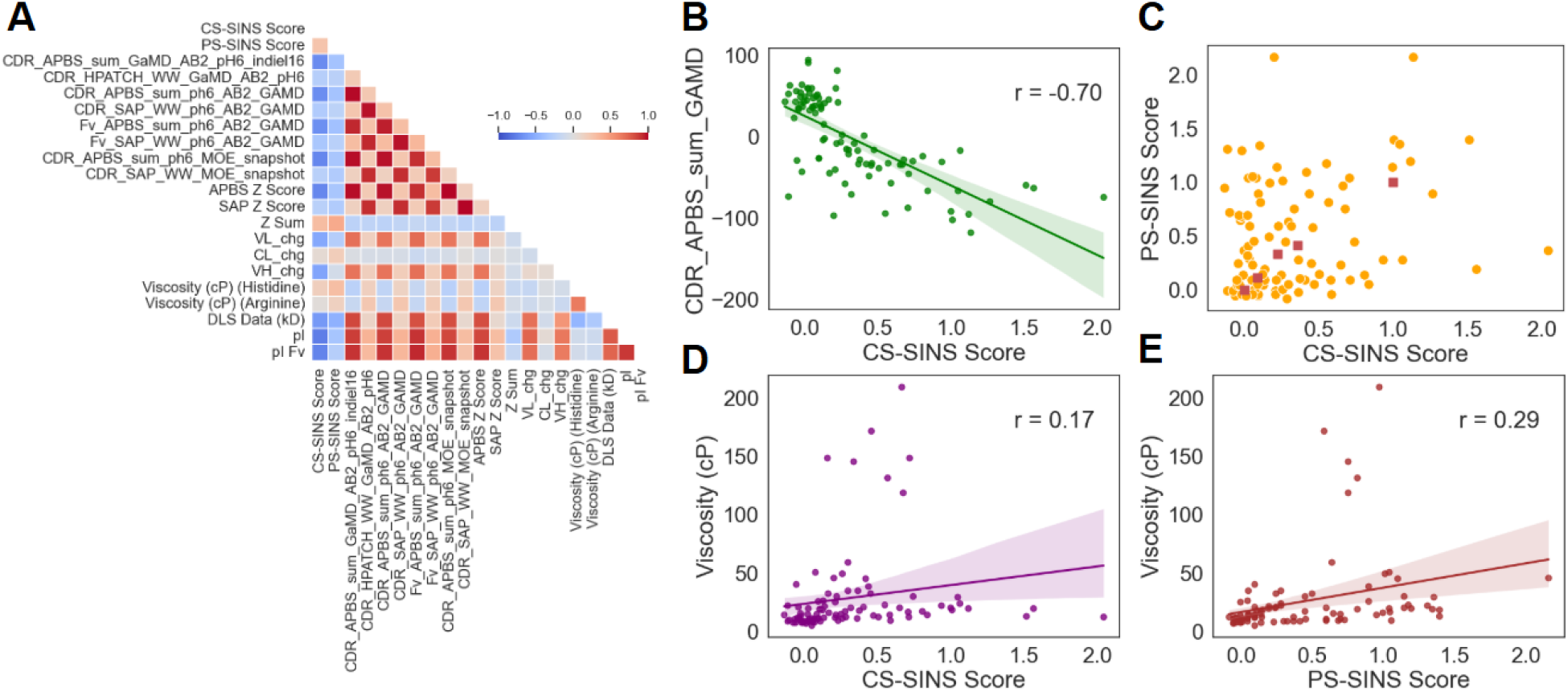
Characterization of SINS and *In Silico* Descriptors. (**A**) The correlation matrix for *in silico* molecular descriptors (described in methods), experimental viscosity data at ∼180 mg/mL) in 20 mM histidine acetate pH 5.8 and 200 mM arginine succinate pH 5.8 buffers, DLS (*k*_D_), and SINS scores (PS-SINS and CS-SINS) for the studied molecules are shown as a heatmap. (**B**) Scatterplot between CS-SINS scores and an *in silico* charge descriptor is shown. (**C**) Relationship between CS-SINS and PS-SINS scores is shown for all the molecules and the five calibration standards are highlighted using square-shaped markers. The dependence of SINS scores on high-concentration viscosity (20 mM histidine acetate pH 5.8 buffer) is shown for - (**D**) CS-SINS and (**E**) PS-SINS. The solid line and shaded area in the scatterplots respectively show the linear regression fit and 95% confidence intervals.

### 2.2. PS-SINS and *in silico* descriptors enable high-throughput prediction of high-concentration viscosity of antibodies

The availability of a large set of antibodies provided with varying viscosity allowed us to evaluate the efficacy of *in silico* and experimental data for prediction of high-concentration viscosity as a binary classification problem. We trained random forest binary classification models^19^ using a leave-one-molecule-out approach with a high-concentration viscosity (∼180 mg/mL, in 20 mM histidine acetate pH 5.8 buffer) cutoff of 15 cP and various combinations of *in silico* descriptors and experimental measurements. The model using *in silico* descriptors of charge and hydrophobicity alone provided only 66% and 71% accuracy, respectively, for predicting the molecules in the low and high viscosity groups (**Figure 3A**). The inclusion of DLS (*k*_D_) and PS-SINS scores along with the *in silico* descriptors improved the classification accuracy of prediction of low and high viscosity molecules to 79% and 86%, respectively (**Figure 3B**). Analysis of the predictor importance scores from the combined model revealed that PS-SINS score and DLS (*k*_D_) are the most influential predictors, underscoring their critical contribution to the improved accuracy (**Figure 3C**). Despite these improvements, a notable number of molecules remained misclassified in both low and high viscosity groups (**Figure 3D**). Examination of the misclassified data revealed a clustering of these molecules near the 15 cP classification threshold. Further inspection of 2D plots combining charge-related descriptors and PS-SINS scores indicated that molecules with negative charge and low PS-SINS scores are particularly prone to misclassification, forming distinct clusters in these plots (**Figure 3E**).

**Figure 3.**
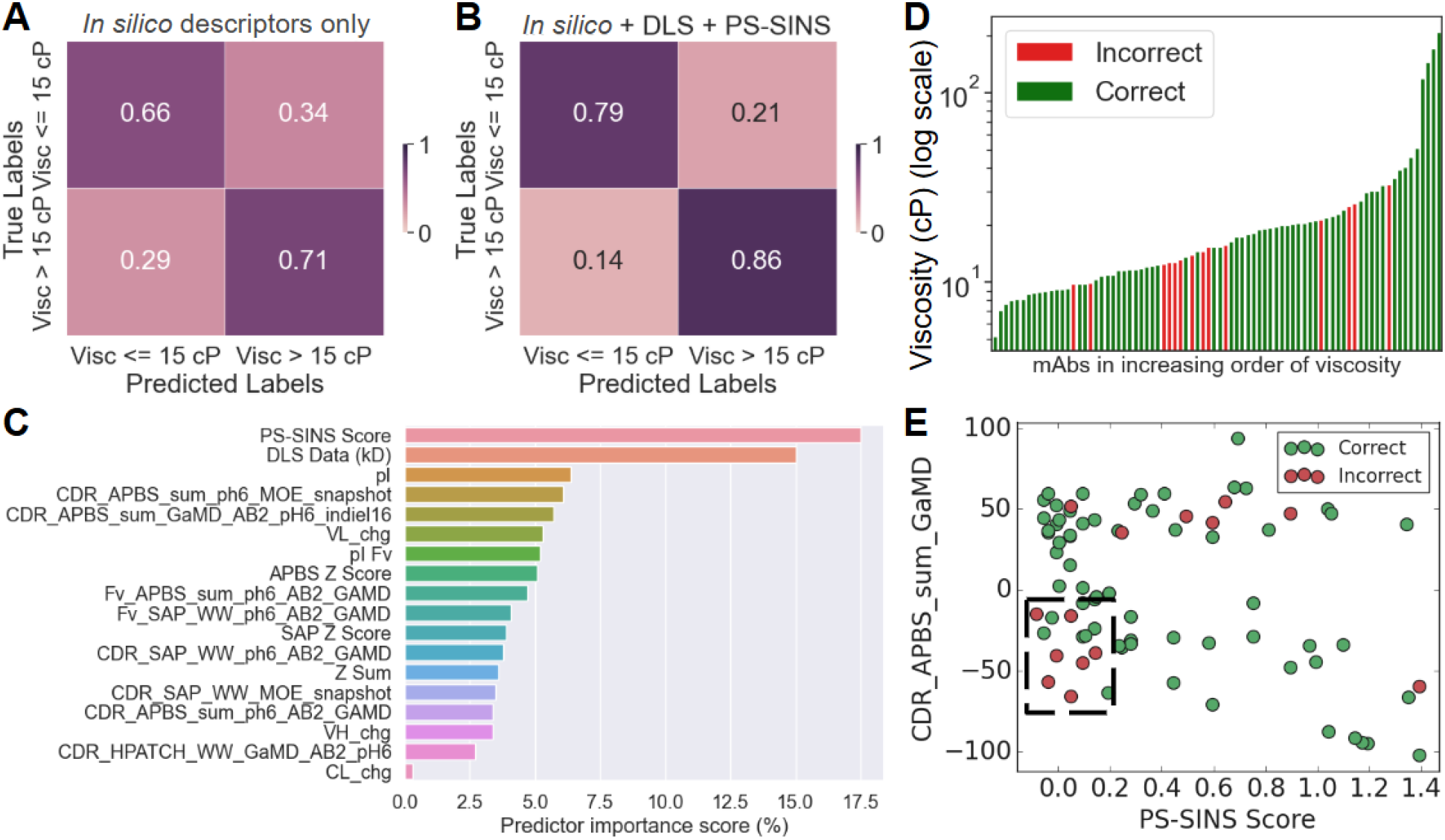
High-throughput Prediction of High-Concentration Viscosity using PS-SINS and *In Silico* Descriptors. (**A**) Binary classification accuracy of a leave-one-sample-out random forest model using only *in silico* descriptors for predicting low and high viscosity (∼180 mg/mL, in 20 mM histidine acetate pH 5.8 buffer) molecules is shown as a confusion matrix. (**B**) Confusion matrix for random forest model with the inclusion of DLS (*k*_D_) and PS-SINS scores alongside *in silico* descriptors is shown. (**C**) Predictor importance scores from the combined model are plotted as a bar plot for the most influential predictors. (**D**) Molecules misclassified by the combined model are shown with respect to their high concentration viscosity values. (**E**) 2D plot combining charge-related descriptors and PS-SINS scores is plotted to show the clustering of misclassified molecules.

### 2.3. PS-SINS in different buffers captures complementary information about the molecule self-association propensity

Motivated by the observed improvement in the random forest prediction accuracy with the incorporation of PS-SINS score in the model, we performed PS-SINS in multiple analysis buffers to evaluate the buffer dependence of the assay and the potential utility of additional buffers in viscosity prediction. The correlation heatmap shows weak correlation between PS-SINS in PBS, histidine-based (10 mM histidine hydrochloride pH 6.0 and 20 mM histidine acetate pH 5.8) and arginine-based (200 mM arginine succinate pH 5.8 and 200 mM arginine chloride pH 5.8) buffers (**Figure 4A**). Furthermore, none of the PS-SINS measurements in various buffers strongly correlate with charge surrogates (CS-SINS, DLS or pI). We incorporated the PS-SINS score in the previous random forest models with PS-SINS measurements from all five buffers and noted high accuracy for high-concentration viscosity prediction despite excluding DLS (*k*_D_) data from the models, which can be cumbersome to obtain during early developability assessment due to low sample availability (**Figure 4B**). Furthermore, we observed that the combination of PS-SINS in just two buffers, PBS and histidine hydrochloride, provides similar overall accuracy without the need for DLS (*k*_D_) measurements (**Figure 4C**).

**Figure 4.**
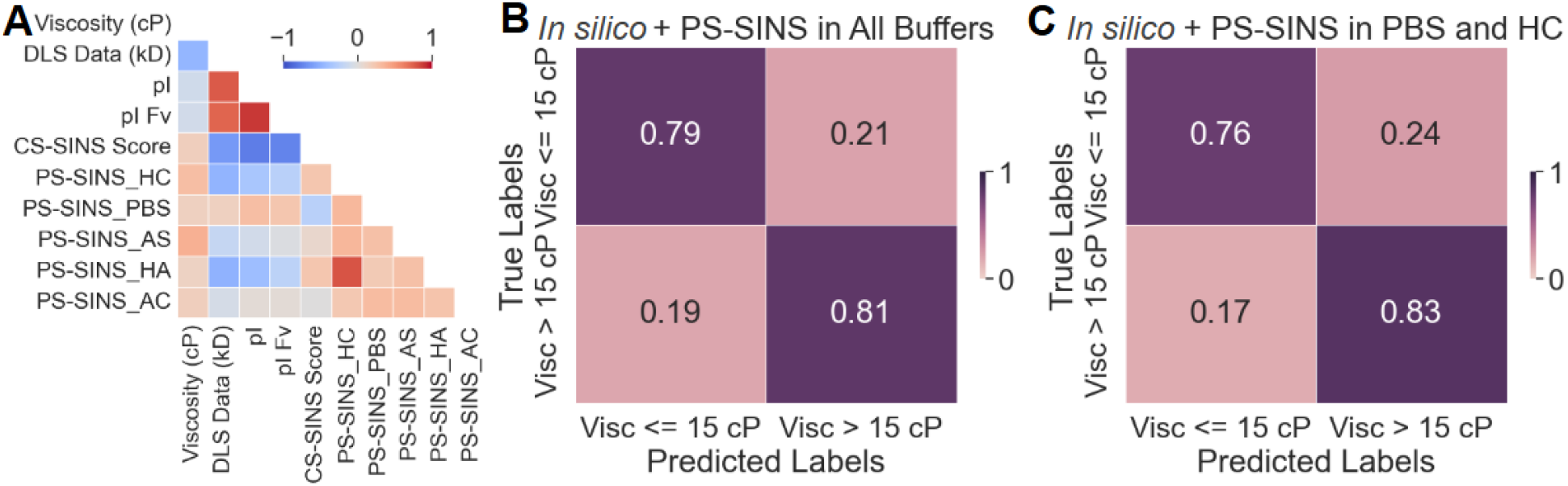
Buffer-Dependent PS-SINS Measurements Capture Complementary Information for Viscosity Prediction. (**A**) Correlation heatmap is shown to capture weak correlations between PS-SINS measurements obtained in different analysis buffers - PBS, 10 mM histidine hydrochloride pH 6.0 (HC), 20 mM histidine acetate pH 5.8 (HA), 200 mM arginine succinate pH 5.8 (AS), and 200 mM arginine chloride pH 5.8 (AC), and their correlations with charge surrogates and high concentration viscosity (∼180 mg/mL, in 20 mM histidine acetate pH 5.8 buffer). Confusion matrix for random forest classification of viscosity using combination of *in silico* descriptors and PS-SINS measurements from (**B**) all five buffers and (**C**) just two buffers (PBS and HC) are shown.

### 2.4. PS-SINS of closely related Fc variants of clinical antibodies suggests putative logarithmic dependence of PS-SINS scores on high-concentration viscosity

While PS-SINS measurements contribute significantly to the multivariate modeling of viscosity, it is not fully clear if viscosity is directly related to PS-SINS score. While all the samples used in this study share the human IgG_1_ heavy chain constant domains and human kappa light chain constant domains, they span a broad range of high-concentration viscosity values. Therefore, to evaluate the direct relationship between PS-SINS scores and high-concentration viscosity, we leveraged the previously characterized Fc variants of two closely related clinically-validated humanized antibodies – trastuzumab and omalizumab – at the low and high ends of the high-concentration viscosity (∼180 mg/mL, in 20 mM histidine acetate pH 5.8 buffer) spectrum.^20^ The Fc variants of omalizumab included the YTE (plasma half-life extension), NG (aglycosylation) and LPLIL (increased effector function) variants, which significantly decreased the viscosity of the parent antibody.^20^ For the trastuzumab, the Fc variants EFT and V12 that result in a significant increase in viscosity have been included among others.^20^ The PS-SINS scores and viscosity measurements for all the Fc variants of both trastuzumab and omalizumab antibodies used in this study are presented in **Tables ST1 and ST2** (Supporting Information), respectively. By plotting the PS-SINS against high-concentration viscosity for both trastuzumab and omalizumab separately, the plots showed weak linear correlation, closely inspecting the distribution of omalizumab variants hinted at the presence of a logarithmic relationship between PS-SINS score and viscosity (**Figure 5A-B**). Therefore, next we plotted both the sets of variants on the same plot and observed a very strong logarithmic relationship between PS-SINS score and viscosity (r = 0.98) (**Figure 5C**). The strong correlation suggests that PS-SINS remains sensitive to the dominant intermolecular forces driving viscosity, even though the anti-Fc capture mechanism could potentially obscure specific Fab–Fc interaction interfaces.

**Figure 5.**
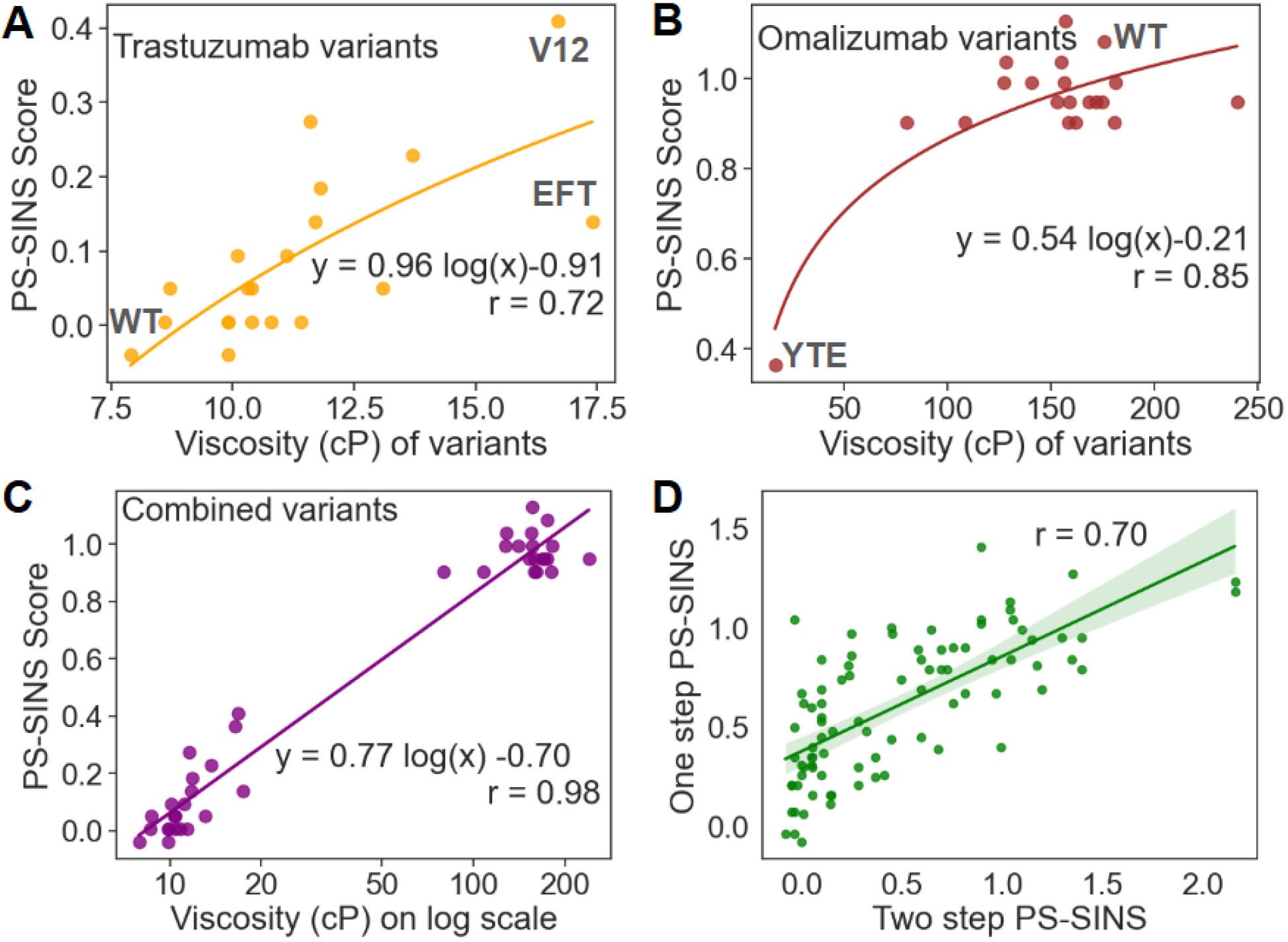
Logarithmic dependence of PS-SINS scores on high-concentration viscosity and performance with one-step purified samples. Scatterplots of PS-SINS score against viscosity for (**A**) trastuzumab and (**B**) omalizumab Fc variants are shown. (**C**) Combined PS-SINS score vs viscosity (log scale for clarity) plot for both sets of Fc variants (trastuzumab and omalizumab) is shown. (**D**) Correlation between PS-SINS scores obtained from one-step purified samples and those obtained from two-step purified samples is shown. The parent (WT) and prominent viscosity altering variants are identified on the plots. The solid lines represent the regression fit line, and the shaded area represents the 95% confidence intervals.

### 2.5. PS-SINS measurements of one-step purified samples correlate with purified material

Finally, we also evaluated the dependence of PS-SINS on the number of protein purification steps for justifying potential use of the assay for developability assessment and early screening of high viscosity variants. We observed that PS-SINS scores obtained after a one-step purification were moderately well correlated (Pearson correlation coefficient, r = 0.70) with those obtained from samples that underwent two steps of purification (**Figure 5D**). We observed that a subset of molecules that had high PS-SINS score for one-step purified material showed lower PS-SINS scores after the second round of purification. This suggests that while using less pure material might lead to the early rejection of some viable candidates (false positives for high viscosity), it is unlikely to erroneously advance high-viscosity candidates to later stages.

## 3. Discussion

This study systematically investigated the potential of emerging Self-Interaction Nanoparticle Spectroscopy (SINS) assays for the early-stage, high-throughput prediction of high-concentration viscosity in monoclonal antibody (mAb) therapeutics. Using a diverse dataset of 96 IgG_1_ antibodies, we provided a comprehensive assessment of both charge-stabilized SINS (CS-SINS) and PEG-stabilized SINS (PS-SINS) assays, critically evaluating their predictive power for high-concentration viscosity and comparing them to *in silico* predictors and traditional biophysical methods. Our findings reveal that CS-SINS scores correlate strongly with charge-related *in silico* molecular descriptors, which aligns with existing literature emphasizing charge as a significant factor in mAb developability.^21,22^ This is mechanistically consistent with the assay’s design, which employs the polycationic polymer poly-L-lysine to stabilize the nanoparticles. The resulting positively charged surface likely biases the interaction landscape, making the assay signal highly sensitive to the electrostatic properties of the test molecule. The strong correlation with charge, coupled with a limited univariate correlation with high-concentration viscosity, suggests that CS-SINS offers restricted additional utility for machine learning-based viscosity prediction models when charge information is already available. Therefore, charge is not the sole determinant of high-concentration viscosity, and other biophysical attributes contribute significantly.^23^

In contrast to the charge-dominated CS-SINS, the PS-SINS assay, when analyzed alone, did not exhibit strong correlations with either charge or hydrophobicity descriptors. When PS-SINS data were integrated with *in silico* descriptors and experimental Dynamic Light Scattering (DLS) diffusion interaction parameter (*k*_D_) data, we observed a significant improvement in the accuracy of random forest-based binary classification models for predicting high-concentration viscosity. These results strongly suggest that while in silico descriptors provide valuable insights into the primary electrostatic and hydrophobic properties of a given molecule, they may not fully account for the complex interactions occurring in a high concentration solution environment. PS-SINS effectively serves as a surrogate for this complex solution-phase behavior, providing orthogonal information that bridges the gap between a given molecule’s calculated surface properties and its macroscopic rheological behavior.^14^

The preliminary success of PS-SINS in a 10 mM histidine hydrochloride (pH 6.0) buffer motivated us to explore the buffer dependence of the assay and the marginal utility of additional buffers in viscosity prediction. Our results showed that PS-SINS measurements in different buffers (PBS, histidine-based, and arginine-based buffers) provided complementary information, as evidenced by their weak cross-correlations. Incorporating PS-SINS scores from multiple buffers into our random forest models significantly improved prediction accuracy for high-concentration viscosity, even in the absence of DLS (*k*_D_) data, which can be cumbersome to obtain early in development due to sample limitations. The inclusion of arginine-based buffers, however, provided negligible classification improvement. As a viscosity-reducing excipient that suppresses protein-protein interactions, arginine results in universally small plasmon wavelength shifts. This compression of the assay’s dynamic range effectively dampens the self-association signals required to differentiate high-from low-viscosity candidates, limiting the predictive utility of PS-SINS in this specific buffer system. Overall, these results suggest that by systematically varying buffer pH, components (e.g., histidine/arginine), and ionic strength (e.g., PBS), the PS-SINS assay can selectively perturb and interrogate different intermolecular forces (e.g., electrostatic and hydrophobic) governing self-association. For instance, a high ionic strength buffer like PBS would shield electrostatic interactions, allowing hydrophobic effects to be more dominant, while a change in pH (as observed previously for omalizumab) would directly modulate the net charge and, consequently, the electrostatic component of self-association.^20,24–26^ This approach, by generating a more comprehensive biophysical fingerprint of a molecule’s developability, underscores the utility of a multi-buffer PS-SINS strategy.

Fc mutations are commonly used to tune pharmacological properties and effector functions, as well as to modulate the intermolecular interactions that govern solution behavior.^27^ Our investigation into Fc mutation variants of clinical antibodies (trastuzumab and omalizumab) provided compelling evidence for a direct and strong logarithmic relationship between PS-SINS score and high-concentration viscosity (r = 0.98), in the context of closely related antibody variants. This correlation remains robust despite the potential for the anti-Fc PS-SINS capture to mask critical Fab-Fc interaction sites. The observed log-linear relationship between high-concentration viscosity and the dilute-solution PS-SINS score is consistent with the established exponential relationship of antibody viscosity with concentration.^10,28^ Since macroscopic viscosity scales exponentially with the magnitude of intermolecular self-association, a linear probe of this tendency—such as the PS-SINS shift—is mathematically expected to correlate with the logarithm of the bulk viscosity. Our findings for the omalizumab and trastuzumab Fc variants powerfully confirm this. The unusually high viscosity of omalizumab is driven by specific Fab-Fab and Fab-Fc interactions.^20,29^ Our Fc variants were designed to directly modulate these interactions and helped demonstrate the sensitivity of PS-SINS assay to such forces, as evinced by strong correlation with the logarithmic changes in viscosity.

Finally, we assessed the impact of antibody purity on PS-SINS assay performance, demonstrating that scores obtained from one-step purified samples correlated reasonably well with those from two-step purified samples. This finding supports the application of PS-SINS at earlier stages of development, where sample purity might be less stringent, enabling even earlier screening and de-risking of promising candidates.^30^ While our study used a diverse set of IgG_1_ molecules, future work could expand the dataset to include other IgG subclasses, non-IgG antibodies, and alternative complex formats to validate the assay’s broad applicability. Similarly splitting such larger datasets into more than two levels of viscosity (e.g. low, medium and high) is expected to yield better classification performance. Further studies could focus on elucidating the precise molecular mechanisms underlying the observed logarithmic relationship between PS-SINS and viscosity. This could involve combining SINS with other biophysical techniques (e.g., small-angle X-ray scattering, molecular dynamics simulations) to gain a more detailed understanding of protein-nanoparticle interactions and their correlation with solution-phase behavior.

In conclusion, this study establishes PS-SINS as a robust high-throughput method for predicting high-concentration viscosity of therapeutic antibodies. The ability to capture orthogonal information, buffer-dependent behavior, direct logarithmic relationship with viscosity coupled with its applicability to less pure samples, position PS-SINS as a powerful tool for accelerating the selection of developable antibody candidates and de-risking the biopharmaceutical development pipeline.

## 4. Methods

### 4.1. PS-SINS

PEG-stabilized self-interaction nanoparticle spectroscopy (PS-SINS) was adapted from the original publication to measure colloidal self-interactions.^14^ Specific (AffiniPure Goat Anti-Human IgG, Fcγ Fragment Specific, Jackson Immunoresearch) and non-specific (ChromPure Goat IgG, Jackson Immunoresearch) capture antibodies were conjugated to 20 nm gold nanoparticles (TedPella). While the use of anti-Fc capture antibodies ensures a consistent orientation of the test molecules on the gold nanoparticles, it is important to note that this tethering may partially mask certain Fab–Fc interaction sites.The resulting conjugates were then stabilized with 2000 MW methoxy PEG thiol (Jenkem Technology). For the analysis, test antibodies were diluted to 1 mg/mL in 10 mM histidine-hydrochloride buffer, pH 6.0. In a 96-well plate, 5 µL of diluted antibody was combined with 15 µL of analysis buffer and 20 µL of the gold conjugate solution. After incubation at room temperature, the absorbance was measured from 501 to 575 nm using a SpectraMax plate reader (Molecular Devices). The plate setup, incubation, and readout were fully automated using a liquid handler (Tecan Fluent). The data was subsequently analyzed using a custom Python script.

### 4.2. CS-SINS

Charge-stabilized self-interaction nanoparticle spectroscopy (CS-SINS) was performed, as previously described, to measure colloidal self-interactions at dilute concentrations.^13^ Anti-Fc gold nanoparticle conjugates were prepared one day prior to the experiment. Briefly, polyclonal goat anti-human Fc IgG (Jackson ImmunoResearch) was buffer-exchanged into a 20 mM sodium acetate buffer (pH 4.3). Similar to PS-SINS, the anti-Fc capture method in CS-SINS could potentially mask some Fab-Fc interactions. This antibody was then mixed with poly-L-lysine and combined with 20 nm gold nanoparticles (Ted Pella) that had been concentrated via centrifugation. For the assay, test monoclonal antibodies were serially diluted to a final concentration of 11.1 µg/mL in the analysis buffer (10 mM histidine chloride, pH 6.0). In a 384-well plate, 5 µL of the nanoparticle conjugate solution was mixed with 45 µL of the diluted antibody solution. The dilution and plate setup have been automated using a liquid handler (Tecan Fluent). After a 4hr incubation at room temperature, the absorbance spectra were measured from 500 to 560 nm using a plate reader (Tecan). The resulting data was processed using a custom Python script.

### 4.3. Sample preparation

#### 4.3.1. Large scale antibody production

The 96 monoclonal antibodies used in this study share the human IgG_1_ heavy chain constant domains and human kappa light chain constant domains. They were selected to span a wide range of charge, hydrophobicity, and high concentration viscosity. The antibodies were generated, as described before, using a high-density transient gene expression (TGE) platform with a CHO K1-derived cell line.^31^ Cells were grown in a seed train to a peak viable cell density of 40−60×10^6^ cells/mL. For transfection, a simplified direct dilution process was employed: cells were not subjected to medium exchange but were instead directly diluted with fresh, proprietary transfection (TFX) medium to a target seeding density of 12×10^6^ cells/mL. A direct transfection was performed by adding a DNA master mix (composed of 2 mg/L coding plasmid, 0.3 mg/L XBP1s plasmid, and filler DNA) and 4.5 mg/L of PEIPro directly to the cell suspension. Cultures were incubated at 33^°^C and 5% CO_2_. Trace elements and valproic acid were added 3–4 hr post-transfection, and additional feeds were supplemented on day 3 and day 7. The supernatant containing the expressed antibody was harvested on day 10 for subsequent purification and analysis. The antibodies were purified using a high-throughput, two-step tandem purification scheme. This method, specifically configured for IgG_1_ mAbs, couples affinity and sizing SEC chromatography systems via three automated AKTA instruments to increase efficiency and decrease resource requirements, aiming for a final product yield of approximately 1 gram per molecule. The tandem method involves two cycles for each step per molecule. For the affinity step (using Purolite Praseto Jetted A50+), a load of 80 g/L was used with 170 mM histidine acetate (pH 2.75) as the elution buffer. This was followed by the size exclusion chromatography (SEC) step (Superdex 200 PG) with up to a 600 mg load in a 15 mL volume.

#### 4.3.2. One-step purified antibody production

1 mL scale IgG expressions were transiently conducted in Expi293F cells (ThermoFisher Scientific). Cells were transfected using 1 mg/mL PEIpro (Cat No: 115-01L). 1 µL PEIpro was diluted into 100 µL serum-free medium, then a 1:1 mixture of heavy chain and light chain plasmids were added and incubated for 10 min. The PEI-DNA complex was added to the 0.85 mL cells, and cells were cultured in a 96-deep well plate at 37 °C and 8% CO_2_ for 7 days, with ∼150 µL fresh medium added 24 hr post transfection. The supernatant was collected and purified with a one-step purification using Protein A affinity resin (MabSelect SuRe™, Cytiva). The absorbance at 280 nm (A_280_) was evaluated to measure protein concentration. Extinction coefficients and isoelectric points (pI) for the antibodies were determined with internal software and confirmed with ExPASy ProtParam tool.^32^

### 4.4. Dynamic light scattering (DLS) measurements

DLS (*k*_D_) measurements were performed on concentrated antibody stock material, formulated in 20 mM histidine acetate pH 5.8 and 200 mM arginine succinate pH 5.8, was filtered using a 0.22 µm PVDF centrifuge filter. Samples were diluted to a concentration range of 2.0 to 10 mg/mL and loaded as triplicates (2 µL) into a STUNNER plate (Unchained Labs) plate using a liquid handling robot (Hamilton). Hydrodynamic radius and UV/Vis data was collected at 25°C and filtered for low deviation (<10%) and high fit quality (sum of squares ≤0.02). The apparent diffusion coefficient (D_app_) was determined, and the diffusion interaction parameter (*k*_D_) was extrapolated via linear regression using the equation D_app_ = D_0_(1 + *k*_D_ *c), where D_0_ and c represent the infinite dilution diffusion coefficient and concentration respectively.^10^ The calculated *k*_D_ quantifies the effect of intermolecular interactions and captures protein self-associative behaviors in solution.

### 4.5. Antibody viscosity measurements

For viscosity measurements, bulk material was aliquoted at 100, 150, and 180 mg/mL and measured using a rheometer (TA instruments DHR30) equipped with a 20 mm 1° cone geometry. Viscosity was measured at 25°C as the 1 minute average at a shear rate of 10^-3^ s^-1^, with the instrument calibrated against S6 and S20 standards.

### 4.6. *In silico* molecular descriptors

A comprehensive suite of *in silico* molecular descriptors was computed for each antibody to characterize their fundamental charge and hydrophobicity properties as described previously.^33^ The charge descriptors, largely derived using the Adaptive Poisson-Boltzmann Solver (APBS), included the sum of electrostatic potential within the complementarity-determining regions (CDRs) at pH 6.0 (CDR_APBS_sum_ph6), and its Z-score normalization (APBS Z Score).^15^ To capture dynamic electrostatic behavior, APBS calculations were also performed on structures derived from Gaussian accelerated molecular dynamics (GaMD) simulations, generating descriptors such as CDR_APBS_sum_GaMD_AB2_pH6_indiel16, CDR_APBS_sum_ph6_AB2_GAMD, and Fv_APBS_sum_ph6_AB2_GAMD, with specific simulation conditions like “AB2” and “indiel16” indicating dielectric constant settings.^17^ Additionally, the net charge of individual antibody domains (e.g., VL_chg, VH_chg, CH1_chg) was calculated based on amino acid composition and protonation states. A static view of CDR electrostatics was provided by CDR_APBS_sum_ph6_MOE_snapshot, derived from Molecular Operating Environment (MOE) simulations.^18^

The hydrophobicity descriptors primarily relied on the Spatial Aggregation Propensity (SAP) method, with CDR_SAP_WW quantifying the aggregation propensity of CDRs based on surface hydrophobicity.^16^ Normalized SAP scores (SAP Z Score) facilitated comparative analysis. Similar to SAP, CDR_HPATCH_WW_GaMD_AB2_pH6 captures hydrophobic patches within CDRs, derived from GaMD simulations. Dynamic hydrophobicity measures from GaMD simulations were also represented by CDR_SAP_WW_ph6_AB2_GAMD and Fv_SAP_WW_ph6_AB2_GAMD for CDRs and the Fv region, respectively. A static assessment of CDR hydrophobicity was obtained from CDR_SAP_WW_MOE_snapshot, an MOE simulation snapshot. These diverse descriptors collectively provided a detailed computational characterization of antibody behavior.

### 4.7. Data Analysis

All data analysis was performed using custom Python scripts using publicly available libraries. For random forest analysis, we used the RandomForestClassifier class in the scikit-learn library.^34^ For leave-one-out analysis, we iteratively left one sample out of training and used it as a test sample. We used oob_score=True, max_features= “sqrt”, class_weight=“balanced_subsample”, and n_estimators = 100 as settings for random forest classification. We used the SciPy library for fitting linear regression lines in the plots.^35^

## Acknowledgements

The authors acknowledge Maniraj Bhagwati, Siok Wan Gan, and Cinzia Stella for the valuable feedback and discussions. Figure 1 has been created using BioRender.

## References

(1) Chan, A. C.; Martyn, G. D.; Carter, P. J. Fifty Years of Monoclonals: The Past, Present and Future of Antibody Therapeutics. Nat. Rev. Immunol. 2025, 25 (10), 745–765. 10.1038/s41577-025-01207-9.

(2) Lu, R.-M.; Hwang, Y.-C.; Liu, I.-J.; Lee, C.-C.; Tsai, H.-Z.; Li, H.-J.; Wu, H.-C. Development of Therapeutic Antibodies for the Treatment of Diseases. J. Biomed. Sci. 2020, 27 (1), 1. 10.1186/s12929-019-0592-z.

(3) Castelli, M. S.; McGonigle, P.; Hornby, P. J. The Pharmacology and Therapeutic Applications of Monoclonal Antibodies. Pharmacol. Res. Perspect. 2019, 7 (6), e00535. 10.1002/prp2.535.

(4) Shire, S. J.; Shahrokh, Z.; Liu, J. Challenges in the Development of High Protein Concentration Formulations. J. Pharm. Sci. 2004, 93 (6), 1390–1402. 10.1002/jps.20079.

(5) Jain, T.; Boland, T.; Vásquez, M. Identifying Developability Risks for Clinical Progression of Antibodies Using High-Throughput in Vitro and in Silico Approaches. mAbs 2023, 15 (1), 2200540. 10.1080/19420862.2023.2200540.

(6) Jezek, J.; Rides, M.; Derham, B.; Moore, J.; Cerasoli, E.; Simler, R.; Perez-Ramirez, B. Viscosity of Concentrated Therapeutic Protein Compositions. Adv. Drug Deliv. Rev. 2011, 63 (13), 1107–1117. 10.1016/j.addr.2011.09.008.

(7) Garidel, P.; Kuhn, A. B.; Schäfer, L. V.; Karow-Zwick, A. R.; Blech, M. High-Concentration Protein Formulations: How High Is High? Eur. J. Pharm. Biopharm. Off. J. Arbeitsgemeinschaft Pharm. Verfahrenstechnik EV 2017, 119, 353–360. 10.1016/j.ejpb.2017.06.029.

(8) Tomar, D. S.; Kumar, S.; Singh, S. K.; Goswami, S.; Li, L. Molecular Basis of High Viscosity in Concentrated Antibody Solutions: Strategies for High Concentration Drug Product Development. mAbs 2016, 8 (2), 216–228. 10.1080/19420862.2015.1128606.

(9) Svilenov, H. L.; Arosio, P.; Menzen, T.; Tessier, P.; Sormanni, P. Approaches to Expand the Conventional Toolbox for Discovery and Selection of Antibodies with Drug-like Physicochemical Properties. mAbs 2023, 15 (1), 2164459. 10.1080/19420862.2022.2164459.

(10) Connolly, B. D.; Petry, C.; Yadav, S.; Demeule, B.; Ciaccio, N.; Moore, J. M. R.; Shire, S. J.; Gokarn, Y. R. Weak Interactions Govern the Viscosity of Concentrated Antibody Solutions: High-Throughput Analysis Using the Diffusion Interaction Parameter. Biophys. J. 2012, 103 (1), 69–78. 10.1016/j.bpj.2012.04.047.

(11) Liu, Y.; Caffry, I.; Wu, J.; Geng, S. B.; Jain, T.; Sun, T.; Reid, F.; Cao, Y.; Estep, P.; Yu, Y.; Vásquez, M.; Tessier, P. M.; Xu, Y. High-Throughput Screening for Developability during Early-Stage Antibody Discovery Using Self-Interaction Nanoparticle Spectroscopy. mAbs 2014, 6 (2), 483–492. 10.4161/mabs.27431.

(12) Wu, J.; Schultz, J. S.; Weldon, C. L.; Sule, S. V.; Chai, Q.; Geng, S. B.; Dickinson, C. D.; Tessier, P. M. Discovery of Highly Soluble Antibodies Prior to Purification Using Affinity-Capture Self-Interaction Nanoparticle Spectroscopy. Protein Eng. Des. Sel. PEDS 2015, 28 (10), 403– 414. 10.1093/protein/gzv045.

(13) Starr, C. G.; Makowski, E. K.; Wu, L.; Berg, B.; Kingsbury, J. S.; Gokarn, Y. R.; Tessier, P. M. Ultradilute Measurements of Self-Association for the Identification of Antibodies with Favorable High-Concentration Solution Properties. Mol. Pharm. 2021, 18 (7), 2744–2753. 10.1021/acs.molpharmaceut.1c00280.

(14) Phan, S.; Walmer, A.; Shaw, E. W.; Chai, Q. High-Throughput Profiling of Antibody Self-Association in Multiple Formulation Conditions by PEG Stabilized Self-Interaction Nanoparticle Spectroscopy. mAbs 14 (1), 2094750. 10.1080/19420862.2022.2094750.

(15) Jurrus, E.; Engel, D.; Star, K.; Monson, K.; Brandi, J.; Felberg, L. E.; Brookes, D. H.; Wilson, L.; Chen, J.; Liles, K.; Chun, M.; Li, P.; Gohara, D. W.; Dolinsky, T.; Konecny, R.; Koes, D. R.; Nielsen, J. E.; Head-Gordon, T.; Geng, W.; Krasny, R.; Wei, G.-W.; Holst, M. J.; McCammon, J. A.; Baker, N. A. Improvements to the APBS Biomolecular Solvation Software Suite. Protein Sci. Publ. Protein Soc. 2018, 27 (1), 112–128. 10.1002/pro.3280.

(16) Chennamsetty, N.; Voynov, V.; Kayser, V.; Helk, B.; Trout, B. L. Design of Therapeutic Proteins with Enhanced Stability. Proc. Natl. Acad. Sci. U. S. A. 2009, 106 (29), 11937–11942. 10.1073/pnas.0904191106.

(17) Miao, Y.; Feher, V. A.; McCammon, J. A. Gaussian Accelerated Molecular Dynamics: Unconstrained Enhanced Sampling and Free Energy Calculation. J. Chem. Theory Comput. 2015, 11 (8), 3584–3595. 10.1021/acs.jctc.5b00436.

(18) Chemical Computing Group Inc. Molecular Operating Environment (MOE). Chem. Comput. Group Inc 2016, 1010.

(19) Breiman, L. Random Forests. Mach. Learn. 2001, 45 (1), 5–32. 10.1023/A:1010933404324.

(20) Heisler, J.; Kovner, D.; Izadi, S.; Zarzar, J.; Carter, P. J. Modulation of the High Concentration Viscosity of IgG1 Antibodies Using Clinically Validated Fc Mutations. mAbs 2024, 16 (1), 2379560. 10.1080/19420862.2024.2379560.

(21) Lehermayr, C.; Mahler, H.-C.; Mäder, K.; Fischer, S. Assessment of Net Charge and Protein-Protein Interactions of Different Monoclonal Antibodies. J. Pharm. Sci. 2011, 100 (7), 2551–2562. 10.1002/jps.22506.

(22) Yadav, S.; Laue, T. M.; Kalonia, D. S.; Singh, S. N.; Shire, S. J. The Influence of Charge Distribution on Self-Association and Viscosity Behavior of Monoclonal Antibody Solutions. Mol. Pharm. 2012, 9 (4), 791–802. 10.1021/mp200566k.

(23) Roberts, C. J. Protein Aggregation and Its Impact on Product Quality. Curr. Opin. Biotechnol. 2014, 30, 211–217. 10.1016/j.copbio.2014.08.001.

(24) Arakawa, T.; Prestrelski, S. J.; Kenney, W. C.; Carpenter, J. F. Factors Affecting Short-Term and Long-Term Stabilities of Proteins. Adv. Drug Deliv. Rev. 2001, 46 (1–3), 307–326. 10.1016/s0169-409x(00)00144-7.

(25) Chen, B.; Bautista, R.; Yu, K.; Zapata, G. A.; Mulkerrin, M. G.; Chamow, S. M. Influence of Histidine on the Stability and Physical Properties of a Fully Human Antibody in Aqueous and Solid Forms. Pharm. Res. 2003, 20 (12), 1952–1960. 10.1023/b:pham.0000008042.15988.c0.

(26) Inoue, N.; Takai, E.; Arakawa, T.; Shiraki, K. Specific Decrease in Solution Viscosity of Antibodies by Arginine for Therapeutic Formulations. Mol. Pharm. 2014, 11 (6), 1889–1896. 10.1021/mp5000218.

(27) Wilkinson, I.; Hale, G. Systematic Analysis of the Varied Designs of 819 Therapeutic Antibodies and Fc Fusion Proteins Assigned International Nonproprietary Names. mAbs 2022, 14 (1), 2123299. 10.1080/19420862.2022.2123299.

(28) Lai, P.-K.; Swan, J. W.; Trout, B. L. Calculation of Therapeutic Antibody Viscosity with Coarse-Grained Models, Hydrodynamic Calculations and Machine Learning-Based Parameters. mAbs 2021, 13 (1), 1907882. 10.1080/19420862.2021.1907882.

(29) Kanai, S.; Liu, J.; Patapoff, T. W.; Shire, S. J. Reversible Self-Association of a Concentrated Monoclonal Antibody Solution Mediated by Fab-Fab Interaction That Impacts Solution Viscosity. J. Pharm. Sci. 2008, 97 (10), 4219–4227. 10.1002/jps.21322.

(30) Chai, Q.; Shih, J.; Weldon, C.; Phan, S.; Jones, B. E. Development of a High-Throughput Solubility Screening Assay for Use in Antibody Discovery. mAbs 2019, 11 (4), 747–756. 10.1080/19420862.2019.1589851.

(31) Wu, R.; Kahl, D. M.; Kloberdanz, R.; Rohilla, K. J.; Balasubramanian, S. Demonstration of a Robust High Cell Density Transient CHO Platform Yielding mAb Titers of up to 2 g/L without Medium Exchange. Biotechnol. Prog. 2024, 40 (3), e3435. 10.1002/btpr.3435.

(32) Gasteiger, E.; Hoogland, C.; Gattiker, A.; Duvaud, S.; Wilkins, M. R.; Appel, R. D.; Bairoch, A. Protein Identification and Analysis Tools on the ExPASy Server. In The Proteomics Protocols Handbook; Walker, J. M., Ed.; Humana Press: Totowa, NJ, 2005; pp 571–607. 10.1385/1-59259-890-0:571.

(33) Park, E.; Izadi, S. Molecular Surface Descriptors to Predict Antibody Developability: Sensitivity to Parameters, Structure Models, and Conformational Sampling. mAbs 2024, 16 (1), 2362788. 10.1080/19420862.2024.2362788.

(34) Pedregosa, F.; Varoquaux, G.; Gramfort, A.; Michel, V.; Thirion, B.; Grisel, O.; Blondel, M.; Prettenhofer, P.; Weiss, R.; Dubourg, V.; Vanderplas, J.; Passos, A.; Cournapeau, D.; Brucher, M.; Perrot, M.; Duchesnay, É. Scikit-Learn: Machine Learning in Python. J Mach Learn Res 2011, 12 (null), 2825–2830.

(35) Virtanen, P.; Gommers, R.; Oliphant, T. E.; Haberland, M.; Reddy, T.; Cournapeau, D.; Burovski, E.; Peterson, P.; Weckesser, W.; Bright, J.; van der Walt, S. J.; Brett, M.; Wilson, J.; Millman, K. J.; Mayorov, N.; Nelson, A. R. J.; Jones, E.; Kern, R.; Larson, E.; Carey, C. J.; Polat, İ.; Feng, Y.; Moore, E. W.; VanderPlas, J.; Laxalde, D.; Perktold, J.; Cimrman, R.; Henriksen, I.; Quintero, E. A.; Harris, C. R.; Archibald, A. M.; Ribeiro, A. H.; Pedregosa, F.; van Mulbregt, P.; SciPy 1.0 Contributors. SciPy 1.0: Fundamental Algorithms for Scientific Computing in Python. Nat. Methods 2020, 17 (3), 261–272. 10.1038/s41592-019-0686-2.

