## Supporting Information for "Self-Interaction Nanoparticle Spectroscopy Predicts High-Concentration Viscosity of Therapeutic IgG1 Antibodies"

**Table ST1. High-concentration viscosity (~180 mg/mL, in 20 mM histidine acetate pH 5.8 buffer) and PS-SINS scores for Fc variants of trastuzumab.**

| Parent | Fc Variant Mutations | Fc Variant Alias | Viscosity (cP)[1] | PS-SINS Score |
|---|---|---|---|---|
| Trastuzumab | None | WT | 8.6 | 0.004 |
| Trastuzumab | HC.M252Y.S254T.T256E | YTE | 9.9 | 0.004 |
| Trastuzumab | HC.L235V.F243L.R292P.Y300L.P396L | VLPLL | 11.6 | 0.274 |
| Trastuzumab | HC.P238D.E233D.G237D.H268D.P271G.A330R | V12 | 16.7 | 0.409 |
| Trastuzumab | HC.S267E.L328F | SELF | 13.7 | 0.229 |
| Trastuzumab | HC.S239D.I332E | SDIE | 11.7 | 0.139 |
| Trastuzumab | HC.S239D.I332E.A330L | SDALIE | 11.1 | 0.094 |
| Trastuzumab | HC.E233P:L234V:L235A:delG236 | PVA# | 10.4 | 0.004 |
| Trastuzumab | HC.N325S.L328F | NSLF | 10.8 | 0.004 |
| Trastuzumab | HC.N297G | NG | 7.9 | -0.040 |
| Trastuzumab | HC.M428L.N434S | MLNS | 10.4 | 0.049 |
| Trastuzumab | HC.F243L.R292P.Y300L.V305I.P396L | LPLIL | 11.8 | 0.184 |
| Trastuzumab | HC.L234A.L235A.P329G | LALAPG | 10.3 | 0.049 |
| Trastuzumab | HC.L234A.L235A | LALA | 8.7 | 0.049 |
| Trastuzumab | HC.K326W.E333S | KWES | 13.1 | 0.049 |
| Trastuzumab | HC1.T366W.HC2.T366S.L368A.Y407V | KiH | 10.1 | 0.094 |
| Trastuzumab | HC.L234F.L235E.D265A | FEA | 9.9 | -0.040 |
| Trastuzumab | HC.S267E.H268F.S324T | EFT | 17.4 | 0.139 |
| Trastuzumab | HC.G236A.S239D.I332E | ADE | 9.9 | 0.004 |
| Trastuzumab | HC.S298A.E333A.K334A | AAA2 | 11.4 | 0.004 |

**Table ST2. High-concentration viscosity (~180 mg/mL, in 20 mM histidine acetate pH 5.8 buffer) and PS-SINS scores for Fc variants of omalizumab.**

| Parent | Fc Variant Mutations | Fc Variant Alias | Viscosity (cP)[1] | PS-SINS Score |
|---|---|---|---|---|
| Omalizumab | None | WT | 175.6 | 1.083 |
| Omalizumab | HC.M252Y.S254T.T256E | YTE | 16.4 | 0.364 |
| Omalizumab | HC.L235V.F243L.R292P.Y300L.P396L | VLPLL | 158.7 | 0.948 |
| Omalizumab | HC.P238D.E233D.G237D.H268D.P271G.A330R | V12 | 156.8 | 1.128 |
| Omalizumab | HC.S267E.L328F | SELF | 180.6 | 0.903 |
| Omalizumab | HC.S239D.I332E | SDIE | 127.1 | 0.993 |
| Omalizumab | HC.S239D.I332E.A330L | SDALIE | 168.0 | 0.948 |
| Omalizumab | HC.E233P:L234V:L235A:delG236 | PVA# | 154.9 | 1.038 |
| Omalizumab | HC.N325S.L328F | NSLF | 174.5 | 0.948 |
| Omalizumab | HC.N297G | NG | 79.8 | 0.903 |
| Omalizumab | HC.M428L.N434S | MLNS | 239.8 | 0.948 |
| Omalizumab | HC.F243L.R292P.Y300L.V305I.P396L | LPLIL | 108.2 | 0.903 |
| Omalizumab | HC.L234A.L235A.P329G | LALAPG | 161.6 | 0.903 |
| Omalizumab | HC.L234A.L235A | LALA | 171.4 | 0.948 |
| Omalizumab | HC.K326W.E333S | KWES | 128.1 | 1.038 |
| Omalizumab | HC1.T366W.HC2.T366S.L368A.Y407V | KiH | 181.2 | 0.993 |
| Omalizumab | HC.L234F.L235E.D265A | FEA | 152.7 | 0.948 |
| Omalizumab | HC.S267E.H268F.S324T | EFT | 156.2 | 0.993 |
| Omalizumab | HC.G236A.S239D.I332E | ADE | 158.3 | 0.903 |
| Omalizumab | HC.S298A.E333A.K334A | AAA2 | 140.3 | 0.993 |

**References**

(1) Heisler, J.; Kovner, D.; Izadi, S.; Zarzar, J.; Carter, P. J. Modulation of the High Concentration Viscosity of IgG1 Antibodies Using Clinically Validated Fc Mutations. *mAbs* **2024**, *16* (1), 2379560. https://doi.org/10.1080/19420862.2024.2379560.
